# The functional differences between paralogous regulators define the control of the General Stress Response in *Sphingopyxis granuli* TFA

**DOI:** 10.1101/2021.10.15.464624

**Authors:** Rubén de Dios, Eduardo Santero, Francisca Reyes-Ramírez

**Author notes:** Authors to whom correspondence should be addressed: R. de Dios and F. Reyes-Ramírez.

## Abstract

*Sphingopyxis granuli* TFA is a contaminant degrading alphaproteobacterium that responds to adverse conditions by inducing the General Stress Response (GSR), an adaptive response that controls the transcription of a variety of genes to overcome adverse conditions. The GSR triggered by TFA is driven by two extracytoplasmic function σ factors (ECFs), EcfG1 and EcfG2, whose functional differences have been addressed previously, being EcfG2 the main activator. Upstream in this cascade, NepR anti-σ factors directly inhibit EcfG activity under non-stress conditions, whereas PhyR response regulators sequester the NepR elements upon stress sensing to relieve EcfG inhibition. These elements, which are essential mediators of the GSR regulation, are duplicated in TFA, being NepR1 and NepR2, and PhyR1 and PhyR2. Here, based on multiple genetic, phenotypical and biochemical evidences including *in vitro* transcription assays, we have assigned distinct functional features to each of these paralogs and assessed their contribution to the GSR regulation, dictating its timing and the intensity. We show that different stress signals are differentially integrated into the GSR by PhyR1 and PhyR2, therefore producing different levels of GSR activation. We demonstrate *in vitro* that both NepR1 and NepR2 bind EcfG1 and EcfG2, although NepR1 produces a more stable interaction than NepR2. Conversely, NepR2 interacts with phosphorylated PhyR1 and PhyR2 more efficiently than NepR1. We propose an integrative model where NepR2 would play a dual negative role: it would directly inhibit the σ factors upon activation of the GSR and it would modulate the GSR activity indirectly by titrating the PhyR regulators.

**IMPORTANCE:** In Alphaproteobacteria, the General Stress Response (GSR) aims at protecting against a variety of stresses. Needing to integrate different signals, its modulation is capital to produce a proportionate response according to the environmental conditions. Individual alphaproteobacterial species have evolved distinct GSR cascades in which the information flow is usually straightforward to ascertain due to the presence of a single copy of at least one of its main regulators (PhyR, NepR and EcfG), restricting the regulatory possibilities. However, *Sphingopyxis granuli* TFA encodes two paralogs of each regulator, multiplying the possible regulatory interplays. We demonstrate that functional differences between paralogous GSR regulators allow an intrinsic feedback regulation in this pathway. We provide evidence of a NepR anti-σ factor that exerts a dual negative feedback regulation on the GSR by interacting with the EcfG σ factors and with the PhyR regulators. This would attune its output to the actual needs of the cell.

## INTRODUCTION

Microbial survivability in natural habitats is usually threatened by fluctuations in the environmental conditions. In order to adapt to these stressing situations, bacteria react by adjusting their transcriptional profile, triggering either specific or global responses, depending on the extent of the transcriptional remodeling. Frequent mechanisms used to control these responses upon exposure to a stimulus are one- or two-component systems, as well as alternative σ factors (Staroń *et al*., 2009). One relevant example of a bacterial global response that is regulated by alternative σ factors is the General Stress Response (GSR), which is a protective broad response that generates cross-protection against a number of unrelated stresses (Staroń & Mascher, 2010). In *Bacillus subtilis* and related Gram-positive bacteria, the GSR is controlled by σ^B^ (Pané-Farré *et al*., 2017), whereas this response is regulated by σ^S^ in many of the proteobacterial representatives of the Gram-negative species (Hengge, 2010; Battesti *et al*., 2011). However, Alphaproteobacteria lack a σ^S^ ortholog (Staroń & Mascher, 2010). In this case, the GSR is regulated by a unique mechanism that combines two-component signalling and transcriptional activation by an extracytoplasmic function σ factor (ECF) (Francez-Charlot *et al*., 2015), which are the most diverse and abundant alternative σ factors (Staroń *et al*., 2009).

In the last decade, the GSR regulatory pathway has been described for a number of alphaproteobacterial representatives (Gourion *et al*., 2009; Bastiat *et al*., 2010; Herrou *et al*., 2012; Jans *et al*., 2013; Kim *et al*., 2013; Fiebig *et al*., 2015; Francez-Charlot *et al*., 2016; Gottschlich *et al*., 2018; Lerdermann *et al*., 2018; Lori *et al*., 2018; Gottschlich *et al*., 2019). The central regulatory elements (the ECF EcfG, its cognate anti-σ factor NepR and the response regulator PhyR) and the mechanistic principles of the signal transduction (Francez-Chralot *et al*., 2009; Campagne *et al*., 2012; Campagne *et al*., 2014) are conserved in most members of this phylogenetic group (Fiebig *et al*., 2015). In the absence of stress, EcfG is sequestered by NepR, preventing the transcription of the GSR regulon (Campagne *et al*., 2012; Herrou *et al*., 2015). Besides, PhyR would remain in its inactive conformation. When a stress appears, it would be sensed by one or more GSR-specific HRXXN histidine kinases, which would phosphorylate PhyR turning it into its active form. In this conformation, PhyR exposes a σ-like domain that is able to interact with NepR more efficiently than its cognate EcfG σ factor, promoting a partner switch (Gourion *et al*., 2008; Francez-Charlot *et al*., 2009; Campagne *et al*., 2012; Herrou *et al*., 2015). This would release EcfG from inhibition, hence activating the transcription of the GSR regulon. Nevertheless, a number of species-specific variations in the signalling circuit may appear (Fiebig *et al*., 2015). Such diversity includes the presence of paralogs of some of the core regulators (Bastiat *et al*., 2010; Staroń & Mascher, 2010; Jans *et al*., 2013; Fiebig *et al*., 2015; Francez-Charlot *et al*., 2015; Francez-Charlot *et al*., 2016), accessory elements involved in the phospho-signalling (Kaczmarczyk *et al*., 2014; Gottschlich *et al*., 2018; Lori *et al*., 2018) or further control at the level of protein stability (Kim *et al*., 2013). Involvement of paralogous regulators is the most common addition to the canonical regulatory pathway. In most cases, the different paralogs display specific functions in the control of the GSR, although with a certain level of redundancy in some instances. For example, in *Sinorhizobium meliloti*, two PhyR homologs (RsiB1 and RsiB2) regulate the GSR to similar extents (Bastiat *et al*., 2010) in response to high temperature and stationary phase. On the other hand, in the same species, the NepR-like anti-σ factors RsiA1 and RsiA2 seemed to control different aspects of the regulation, since the deletion of *rsiA2* led to derepression of the response, whereas *rsiA1* mutation resulted in lethality (Bastiat *et al*., 2010). The most accentuated known example of GSR regulator multiplicity is found in *Methylobacterium extorquens*, in which up to six EcfG paralogs are involved in the control of the response, with EcfG1 and EcfG2 playing a major role in the stress resistance (Francez-Charlot *et al*., 2016). Furthermore, a main NepR protein seem to play a canonical anti-σ role, inhibiting two EcfG paralogs (EcfG1 and EcfG5 to a certain extent) and being amenable to PhyR sequestration, whereas an additional NepR copy (MexAM1_META2p0735) is unable to interact with any of the EcfG paralogs. Rather, it interacts with PhyR and produces a negative effect on the GSR activity, thus suggesting it would act as an anti-anti-anti-σ factor, which implies a divergent functional role in the regulation with respect to the main NepR paralog (Francez-Charlot *et al*., 2016). Moreover, a similar NepR paralog specialization has also been proposed in *Sphingomonas melonis* for NepR2 with respect to NepR (Gottschlich *et al*., 2019).

*Sphingopyxis granuli* TFA is an alphaproteobacterium that has been deeply characterized regarding its ability to use the organic solvent tetralin as carbon and energy source, both at the biochemical and genetic level (reviewed in Floriano *et al*., 2019). Also, since the annotation of its genome and after confirmation by functional characterization (García-Romero *et al*., 2016, it has been defined as the first facultative anaerobe within the *Sphingopyxis* genus due to its capability to respire nitrate anaerobically, and its global regulatory response to this condition has been described (González-Flores *et al*., 2019; González-Flores *et al*., 2020). Recently, the GSR regulators encoded in TFA were identified (de Dios *et al*., 2020). This strain encodes two paralogs of each of the regulators of the central GSR pathway, distributed in two genomic loci: one bearing *nepR1* and *phyR1*, and other genomic location containing *nepR2* and *ecfG1* in a bicistronic operon, *ecfG2* and *phyR2*. The individual roles of EcfG1 and EcfG2 in the regulation have been investigated (de Dios *et al*., 2020), being EcfG2 the main GSR activator, as it confers stress resistance by itself and is able to control the expression of the whole GSR regulon. On the other hand, EcfG1 seems to play an accessory role, since its expression is EcfG2-dependent and it is only able to fully activate the transcription of part of the GSR target genes.

In this work we have further characterized the GSR regulatory pathway in TFA by combining *in vivo* and *in vitro* approaches. We show a functional differentiation between NepR1 and NepR2 in the control of the response and a different specificity in the stress signalling by PhyR1 and PhyR2. Finally, after reproducing the regulatory system *in vitro*, we propose an integrative model in which the PhyR regulators would produce different levels of activation of the GSR according to the stress that triggers it. Also, in this model NepR2 would play a dual role: it would directly inhibit the EcfG σ factors and it would negatively modulate the GSR activity indirectly, by titrating the PhyR regulators and releasing NepR1 to further inhibit EcfG1 and EcfG2, thus preventing an overactivation of the response.

## RESULTS

### 1. NepR1 and NepR2 play specific roles in the regulation of the GSR

Previous analysis of the TFA genome annotation revealed that the elements involved in the core GSR signalling pathway appear duplicated (de Dios 2020). EcfG1 and EcfG2 are the σ factors that drive the transcription of the GSR regulon, with EcfG2 having the leading role in the activation (de Dios *et al*., 2020). Upstream in the signalling cascade, the NepR1 and NepR2 paralogs would act as anti-σ factors, inhibiting the GSR in the absence of stress. In the genome, *nepR1* is transcribed in a monocistronic operon, presenting up to two suboptimal GSR target promoters upstream its coding region (Sup. Fig. S1A). This is coherent with a subtle increase in transcription under GSR-inducing conditions, according to differential RNA-seq (dRNA-seq) data and RT-qPCR (Sup. Fig. S1B). In contrast, *nepR2* is transcribed as the first gene in the *nepR2ecfG1* operon in a GSR-dependent manner, presenting a canonical GSR target promoter upstream (Sup. Fig. S1A) (de Dios *et al*., 2020). This causes a strong upregulation of *nepR2* transcription under GSR-inducing conditions, as shown by previous dRNA-seq data (de Dios *et al*., 2020) and RT-qPCR measurements (Sup. Fig. S1B). Due to their inhibitory function, their absence would theoretically lead to a derepression of the response under non-stress conditions.

In order to address their role in the regulation, the construction of the different *nepR* deletion mutants was attempted. However, in other Alphaproteobacteria (Bastiat *et al*., 2010; Lourenço *et al*., 2011), the deletion of a *nepR* homolog that is co-transcribed together with an EcfG coding gene in an autoregulated operon resulted in lethality. This has been argued to be due to an uncontrolled transcriptional activity of the respective EcfG ortholog on its own promoter in the absence of NepR, which may lead to a deleterious overactivation of the GSR. In agreement with this, *nepR1* could be deleted in TFA, contrarily to *nepR2*. Nevertheless, a deletion mutant in the whole *nepR2ecfG1* operon could be constructed. To address the cause of the *nepR2* essentiality, *in trans* complementation experiments were performed. In these assays, the viability of the Δ*nepR2ecfG1* mutant was assessed after transformation with a plasmid bearing *ecfG1* without the promoter region, preceded by its own promoter or by a GSR-insensitive promoter. As shown in Sup. Fig. S2, the plasmid bearing *ecfG1* under its own promoter was the only one unable to be stabilized in the mutant, which highlights the essentiality of NepR2 to control the autoinduction of *ecfG1*.

To distinguish the specific role of each NepR paralog in the GSR regulation, a *nepR2::lacZ* reporter (which has been previously used to assess the GSR activity in TFA (de Dios *et al*., 2020)) was integrated in the chromosome of the Δ*nepR1* and the Δ*nepR2ecfG1* mutants, as well as in the Δ*nepR1*Δ*nepR2ecfG1* triple mutant. Next, their β-galactosidase activity was measured in exponential (GSR repressed) and stationary phase (GSR active) and compared to those of the wild type and the Δ*ecfG1* single mutant (the timepoints of activity measurement are specified in Sup. Fig. S3). According to the results shown in Fig. 1, the Δ*ecfG1* mutant showed a slightly lower level of GSR activity in stationary phase (as previously reported in de Dios *et al*., 2020), whilst the Δ*nepR1* mutant presented a derepressed GSR in exponential phase compared to the wild type, with a slight increase in stationary phase. The Δ*nepR2ecfG1* performed similarly to the wild type and the Δ*ecfG1* mutant in the repression of the GSR under exponential growth. However, in stationary phase, this mutant nearly doubled the activity of the wild type strain. In the case of the Δ*nepR1*Δ*nepR2ecfG1* mutant, in which a constitutively active EcfG2 would be alone to activate the response, a strong derepression was observed in exponential phase, presenting approximately a 40-fold increase in activity compared to the wild type TFA in exponential phase, which continued to increase in stationary phase to even higher levels. The levels of activity reached by the triple mutant indicate that, even under stress conditions, the maximum levels of GSR expression are not reached by the wild type strain, suggesting that a proportion of the anti-σ factors remain active under our experimental conditions. Altogether, these results suggest that NepR1 and NepR2 have specifics roles in GSR regulation, with NepR1 playing a main role in the global repression of the GSR in TFA in the absence of stress, and hence in its initial activation, and with NepR2 modulating the intensity of the response once it is active.

**Figure 1.**
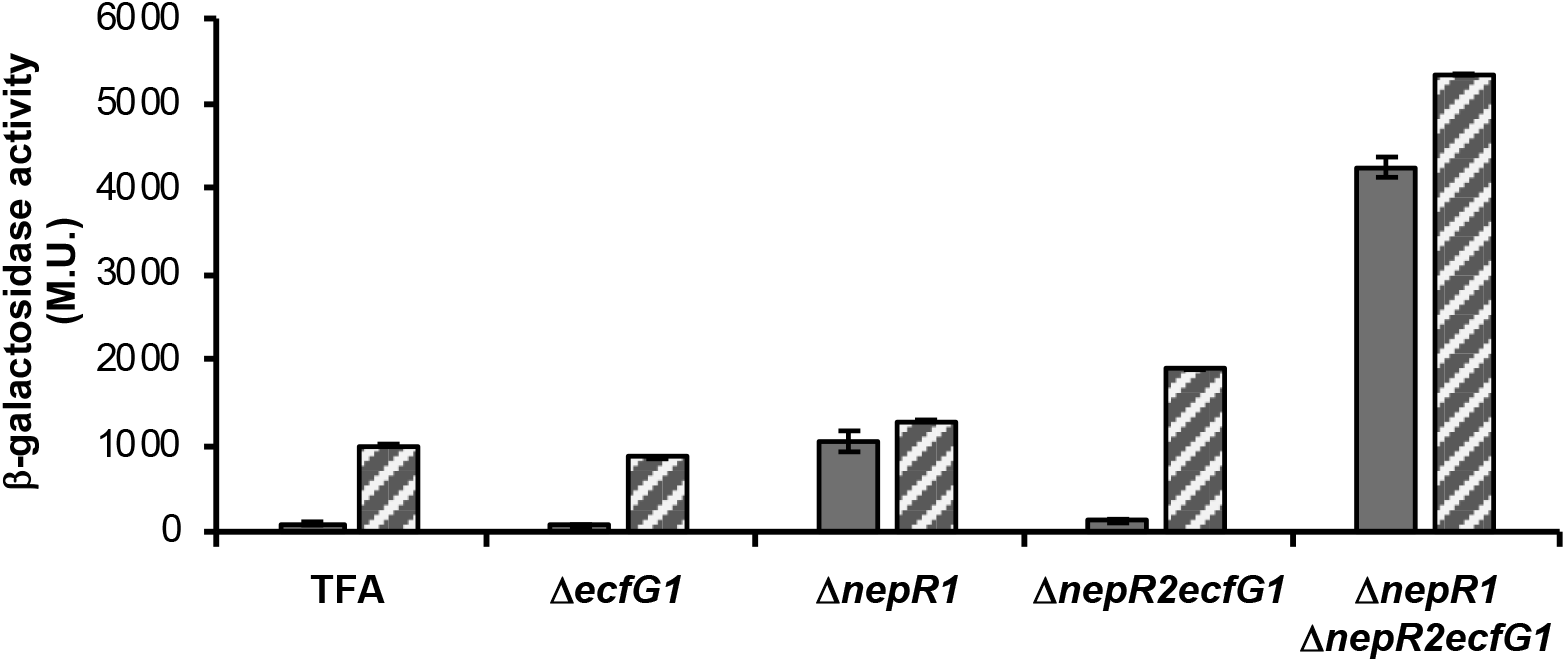
β-galactosidase activity from the *nepR2::lacZ* translational fusion in different Δ*nepR* mutant backgrounds compared to the wild type and the Δ*ecfG1* single mutant. The activity was measured in exponential (whole bars) and stationary phase (striped bars).

### 2. NepR1 and NepR2 show different binding affinities for EcfG1 and EcfG2

Structural studies describing the molecular aspects of the partner switching mechanism that mediate the GSR activation in Alphaproteobacteria revealed that the EcfG inhibition by NepR occurs by a direct protein-protein interaction (Campagne *et al*., 2012). Since TFA encodes two paralogs of each of these proteins, one possible model would be that each of the EcfG proteins were specifically titrated by one of the NepR anti-σ factors. To explore this option, combinatory mutants were constructed (namely, Δ*nepR1*Δ*ecfG1*, Δ*nepR1*Δ*ecfG2* double mutants and Δ*nepR1*Δ*ecfG1*Δe*cfG2* triple mutant) and their ability to resist to heavy metals and osmotic stress were tested. As a result (Sup. Fig. S4A) only those strains lacking *ecfG2* showed an increased sensitivity compared to the wild type. Contrarily, when β-galactosidase activity from the *nepR2::lacZ* fusion was measured in those backgrounds, it reached higher levels in the Δ*nepR1*Δ*ecfG2* mutant compared to those of the Δ*nepR1*Δ*ecfG1* (even beyond those of the wild type TFA) as shown in Sup. Fig. S4B. These results imply that each EcfG paralog is not specifically titrated by one NepR protein. Rather, they would suggest a more complex interplay at the NepR-EcfG interface, which may be defined by the protein-protein affinities between each of the σ-anti-σ pairs and the relative abundance of these regulators in the cell. In order to characterise the four possible NepR-EcfG interactions (NepR1 with EcfG1 or EcfG2 and NepR2 with EcfG1 or EcfG2) and their effect on the transcriptional output of the response, each of these regulators was purified. After that, they were used in different combinations in an *in vitro* transcription (IVT) setup together with the native core RNAP purified from TFA and using the *P*_*nepR2*_ promoter as template (de Dios *et al*., 2020). After fixing a common concentration for each EcfG paralog below RNAP saturation levels (de Dios *et al*., 2020), either NepR1 or NepR2 were added to the reactions in increasing molecular proportion with respect to them (Fig. 2A). As a result, NepR1 was able to titrate either EcfG protein nearly in a 1:1 proportion, achieving a complete inhibition of transcription. In contrast, a 10:1 molecular excess of NepR2 with respect to either EcfG1 or EcfG2 could not reach similar levels of inhibition to those of NepR1, indicating a weaker interaction between NepR2 and the EcfG σ factors compared to that of NepR1. To further address this interplay, the NepR-EcfG protein-protein interactions were quantified by surface plasmon resonance. For these experiments either NepR1 or NepR2 were immobilised on CM5 chip and either EcfG1 or EcfG2 were injected as analytes under a continuous flow. Kinetic analysis of the interactions for each NepR-EcfG pair gave the respective dissociation constants (K_D_) shown in Fig. 2A. These results agree with those obtained with the *in vitro* transcription system, showing a correlation between lower K_D_ values and stronger repression of gene transcription. Thus, a stronger interaction between those EcfG-NepR pairs including NepR1 would be responsible for a more efficient repression of transcription compared to those pairs including NepR2.

**Figure 2.**
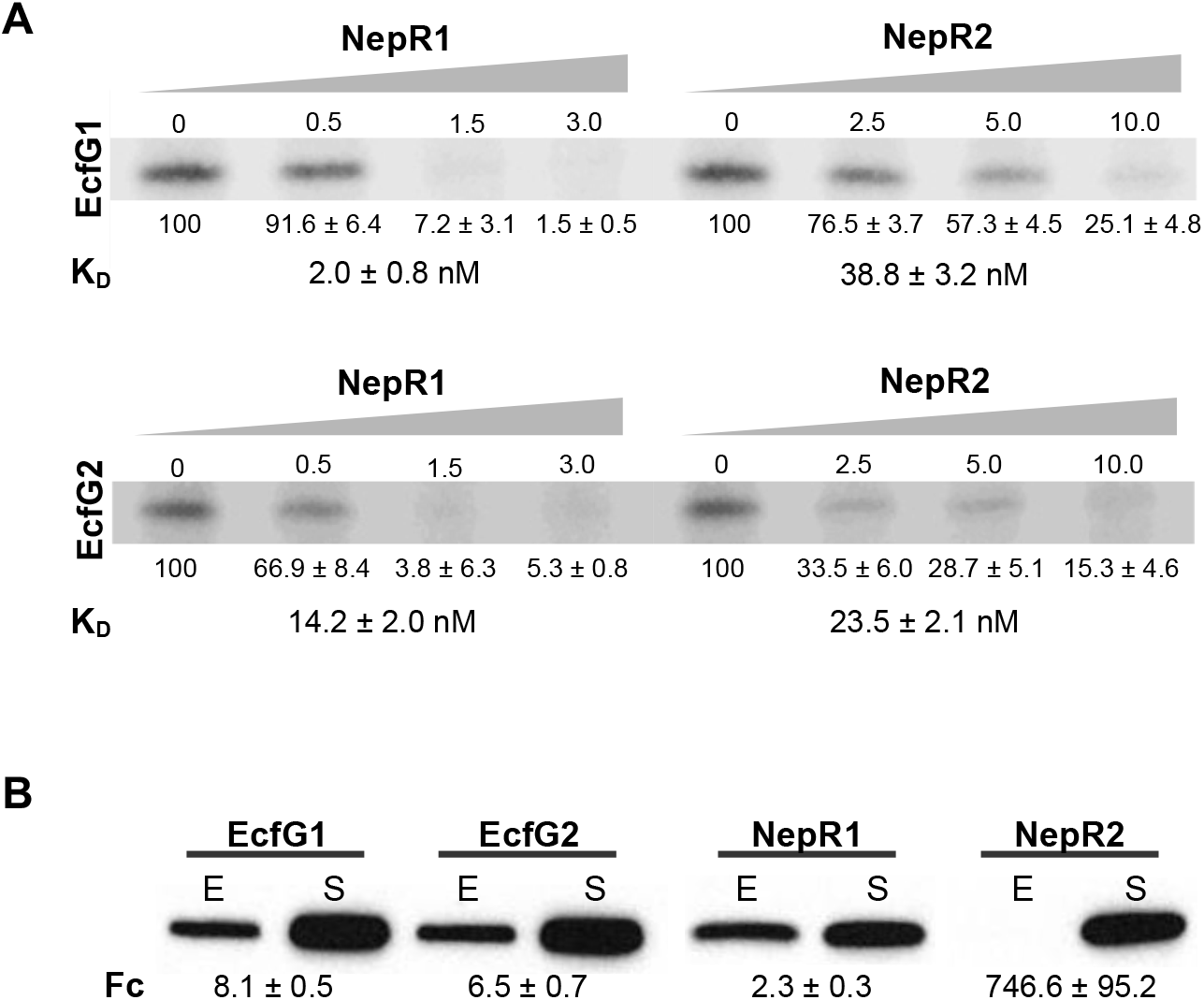
*In vitro* transcription levels defined by the interaction between the different EcfG-NepR pairs encoded in TFA and protein quantification of the different regulators. A) IVT results using either EcfG1 or EcfG2 as σ factor and increasing concentrations of either NepR1 or NepR2. Transcription quantifications are referred to those obtained in the absence of anti-σ factor. The dissociation constant (K_D_) measured for each EcfG-NepR pair using surface plasmon resonance is indicated underneath each combination. B) Immunodetection of EcfG1, EcfG2 NepR1 and NepR2 tagged in their C-terminal end with a 3xFLAG epitope. Samples were collected in exponential (E) and stationary phase (S). Protein accumulation fold-change (Fc) is indicated underneath.

Apart from the affinity between the different NepR-EcfG pairs, the relative amounts of each of the elements involved in an interaction also determines its output. To have an impression of the evolution of the *in vivo* protein accumulation of each of the NepR and EcfG regulators, FLAG-tagged versions of each of them were constructed in a wild type background. Their accumulation was assessed by Western blot in exponential phase (in which the GSR would be off due to NepR inhibition) and in stationary phase (in which the GSR is active because of prevention of the NepR-EcfG interaction). As a result, a general increase in the accumulation of the four regulators was observed in stationary phase, with the most drastic change being that of NepR2 (Fig. 2B). These results are coherent with those obtained in the *in vitro* transcription assays, since NepR2 would be needed in bigger amounts than NepR1 in order to perform an efficient inhibition.

### 3. The GSR is specifically activated by PhyR1, PhyR2 or both of them depending on the stress

The role of PhyR response regulators consists in derepressing the GSR upon receiving the stress signal in the shape of phosphorylation by sequestering NepR proteins, thus acting as indirect activators of the GSR regulon. In other alphaproteobacterial species, *phyR* mutants behave similarly to *ecfG* mutants regarding their stress resistance, displaying an increased sensitivity compared to the parental wild type strain. This is due to the inability of these strains to prevent EcfG titration by NepR.

In order to address the role of each PhyR paralog encoded in TFA in the GSR signalling, deletion mutants were constructed in each *phyR* gene, as well as a double mutant. Subsequently, the resulting mutant strains were challenged to resist a variety of stresses compared to the wild type strain and a Δ*ecfG1*Δ*ecfG2* double mutant, which is totally impaired in the GSR activation. The results revealed that an increased sensitivity to heavy metals (copper) was only observed in those Δ*phyR* mutant backgrounds lacking *phyR2* (Fig. 3A). On the other hand, an increased sensitivity to oxidative stress was obtained only in the absence of *phyR1* (Fig. 3B). Regarding the resistance to desiccation, all Δ*phyR* mutant strains were affected compared to the wild type, with a milder sensitivity observed for the Δ*phyR2* mutant (Fig. 3C). In contrast, only the Δ*phyR1*Δ*phyR2* double mutant resulted more affected than the wild type under osmotic stress conditions (Fig. 3A). Altogether, this suggests that PhyR1 and PhyR2 are activated specifically depending on the stress that triggers the GSR signalling.

**Figure 3.**
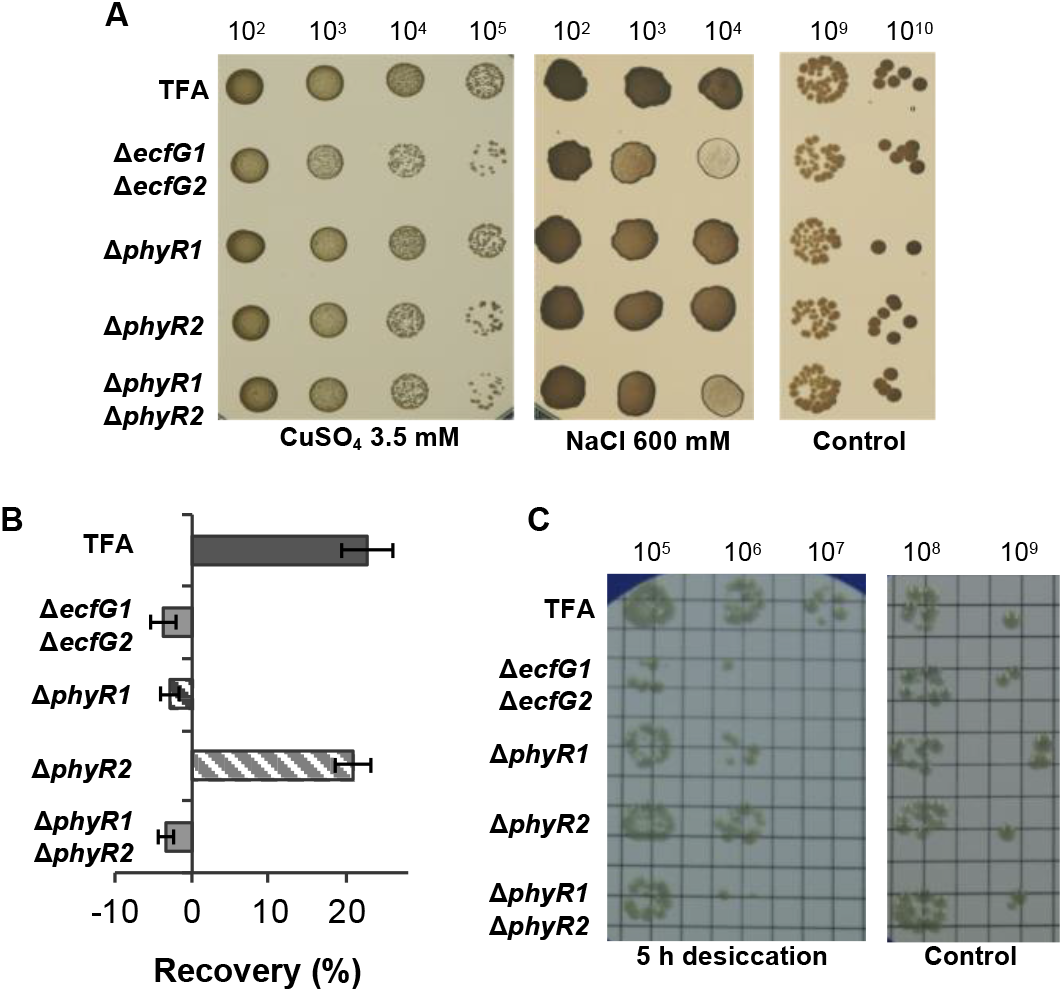
Stress resistance phenotypes of the Δ*phyR1* and Δ*phyR2* single mutants and the Δ*phyR1*Δ*phyR2* double mutant compared to the wild type TFA and the Δ*ecfG1*Δ*ecfG2* double mutant (stress-sensitive control). The phenotypes tested were A) resistance to CuSO_4_ 3.5 mM and NaCl 600 mM, B) exposure to desiccation during 5 h and C) recovery of the growth after the addition of H_2_O_2_ 10 mM.

### 4. PhyR1 and PhyR2 produce different levels of activation of the GSR

The results presented previously conveyed the idea that each of the PhyR regulators encoded in TFA performed distinctive roles in the GSR activation. To evaluate their ability to activate the response, the *nepR2::lacZ* reporter was introduced in each of the Δ*phyR* mutant backgrounds and their β-galactosidase activity was measured in exponential and stationary phase compared to that of the wild type (Fig. 4). As expected, the Δ*phyR1*Δ*phyR2* mutant showed a similar level of activity to that of the Δ*ecfG1*Δ*ecfG2* mutant. The Δ*phyR1* single mutant showed a marked decrease in the activity, mainly observed in stationary phase, whereas the Δ*phyR2* mutant produced slightly lower levels of activity than the wild type. These results indicate that PhyR1 is able to produce a stronger activation of the GSR than PhyR2, at least in stationary phase induced by carbon starvation.

**Figure 4.**
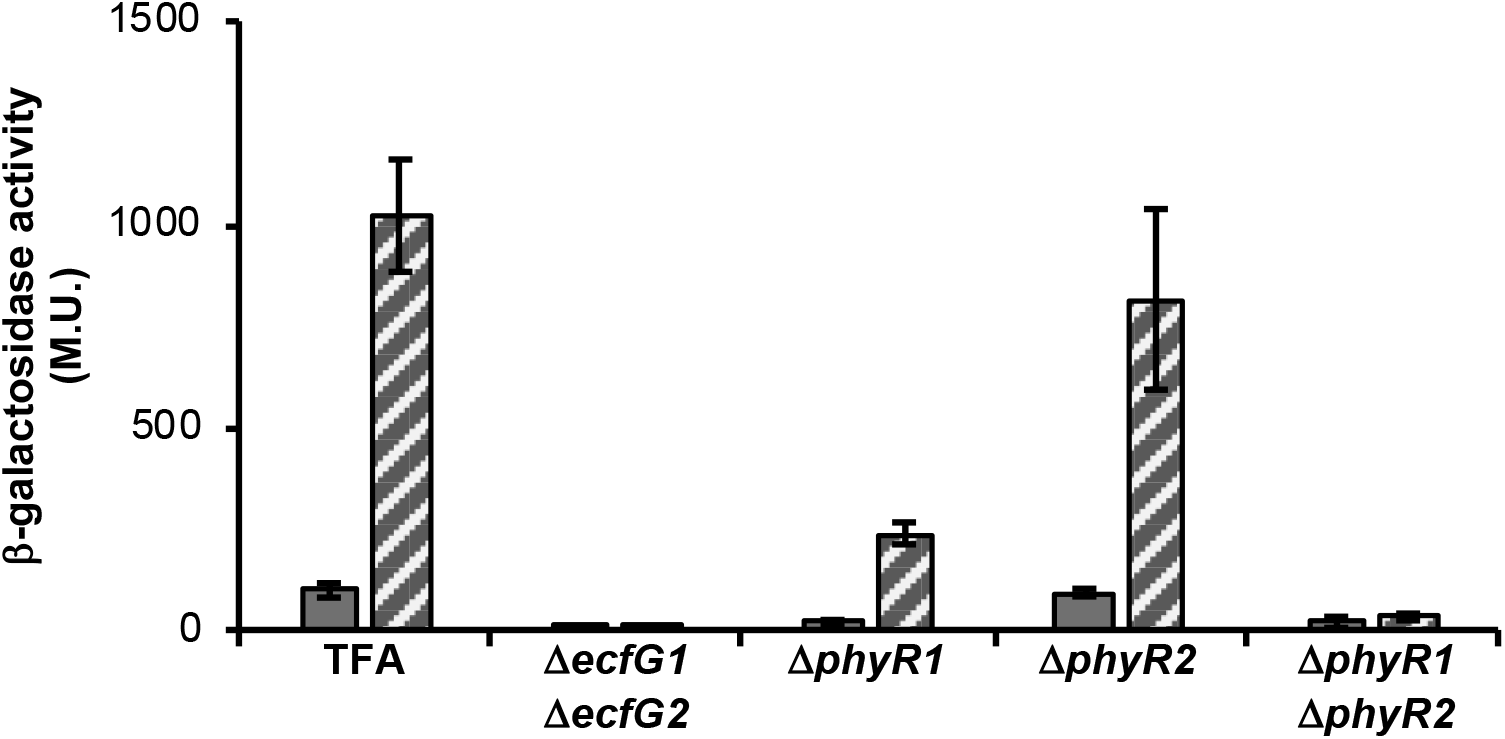
β-galactosidase activity from the *nepR2::lacZ* translational fusion in different Δ*phyR* mutant backgrounds compared to the wild type and the Δ*ecfG1*Δ*ecfG2* double mutant (negative control). The activity was measured in exponential (whole bars) and stationary phase (striped bars).

After comparing β-galactosidase activity from the *nepR2::lacZ* fusion in the different Δ*phyR* mutants to those of the Δ*ecfG* mutants (de Dios *et al*., 2020), similarities in both expression patterns were observed (i. e, the expression phenotype of the Δ*phyR1* mutant resembled that of an Δ*ecfG2* mutant, and the phenotype of the Δ*phyR2* mutant resembled that of the Δ*ecfG1*). This raised the question whether there would be a specific signalling from PhyR1 toward EcfG2 and from PhyR2 toward EcfG1. Nevertheless, a Δ*phyR1*Δ*ecfG1* double mutant, in which the only signalling stream possible would be from PhyR2 to EcfG2, showed a similar expression to that observed in the Δ*phyR1* single mutant (Sup. Fig. S5). This suggests that PhyR2, as well as PhyR1, are able to communicate stress to EcfG2, opening the possibility of a signal convergence via the NepR anti-σ factors.

### 5. PhyR1 and PhyR2 are able to interact more efficiently with NepR2 than with NepR1

As demonstrated for other alphaproteobacterial species, the only stream that the GSR signalling pathway follows is the PhyR-NepR-EcfG cascade, with no accessory regulation occurring between the PhyR and EcfG regulators known so far. Therefore, the only possibility that PhyR1 or PhyR2 may have to activate the transcription would be the direct interaction with either NepR1 or NepR2 in a 1:1 theoretical proportion.

In order to determine the ability of PhyR1 and PhyR2 to activate the GSR, they were purified and added to the previously set up IVT system. All possible PhyR-NepR-EcfG combinations were assayed, using a molecular NepR-EcfG proportion that would *a priori* inhibit transcription, such as 1.5:1 for the NepR1-EcfG pairs and 10:1 for the NepR2-EcfG pairs. The PhyR ratio used in the assays were 2:1 with respect to NepR1 and 1:1 with respect to NepR2. To simulate an active or inactive status of the GSR, defined by the phosphorylation state of the PhyR proteins, the universal phosphor-donor acetyl phosphate (or a mock treatment) was added to the reactions accordingly. The results (Fig. 5, with an extended version presented in Sup. Fig. S6) show that only phosphorylated PhyR1 and PhyR2 were able to stimulate transcription using either EcfG1 or EcfG2. Therefore, when acetyl phosphate was not added, transcription levels remained insensitive to the presence of either PhyR1 or PhyR2. Regarding the anti-σ factor used in each case, whereas both active PhyR1 and PhyR2 could relieve the inhibition exerted by NepR2 to different extents, only PhyR1 was able to activate transcription *in vitro* to detectable levels in the presence of NepR1 in the conditions tested (6.1-fold for PhyR1 versus 1.3-fold for PhR2 using EcfG1 as σ factor; 1.2-fold for PhyR1 and no transcription stimulation by PhyR2 when adding EcfG2). This is coherent with the β-galactosidase activity results obtained using the *nepR2::lacZ* reporter, (814.6 M.U. in the Δ*phyR2* mutant versus 237.8 M.U. in the Δ*phyR1*, as shown in Fig. 4) thus confirming the greater potential of PhyR1 to trigger the GSR compared to PhyR2.

**Figure 5.**
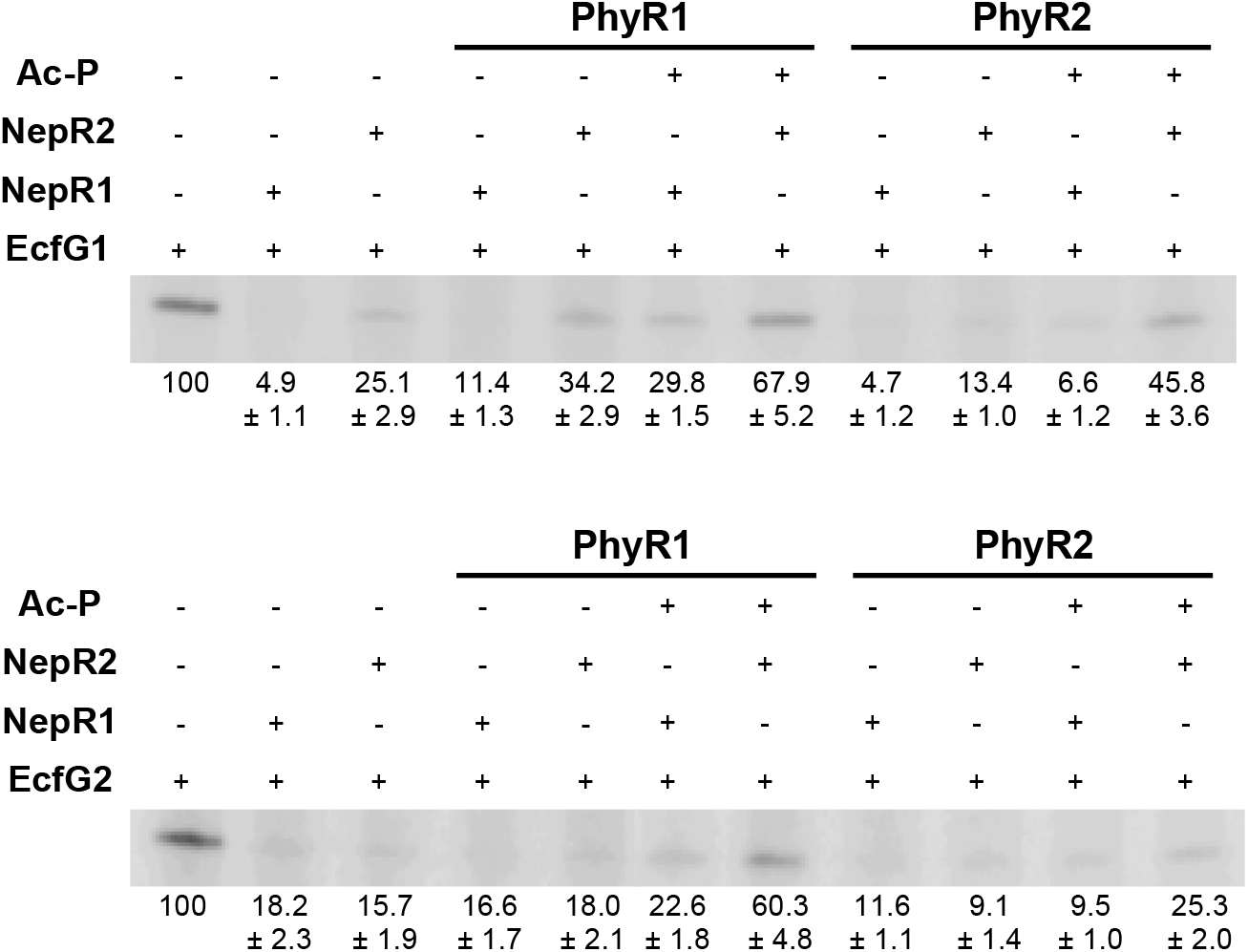
*In vitro* reconstruction of the GSR using 0.2 μM of either EcfG1 or EcfG2 as σ factor in an *in vitro* transcription system. The molecular proportions among all proteins added to each reaction (EcfG:NepR:PhyR) were 1:1.5:3 when using NepR1 as anti-σ factor and 1:10:10 when using NepR2. When required, 15 mM acetyl phosphate (AcP) was added to obtain phosphorylated versions of the PhyR proteins. Transcription quantifications are referred to those obtained in the absence of NepR and PhyR proteins.

An intriguing observation from this data is the higher transcription levels obtained when using NepR2 in the presence of any of the phosphorylated PhyR proteins than when using NepR1. In the context of the dynamic protein-protein interactions that regulate the alphaproteobacterial GSR, this would mean that NepR1 is able to interact more efficiently with the EcfG σ factors than with the PhyR proteins (regardless of their phosphorylation state), contrarily to NepR2, which would present higher affinity for the active PhyR proteins than for the σ factors.

## DISCUSSION

*S. granuli* TFA is an alphaproteobacterium that encodes two paralogs of each of the central regulators of the GSR. Paralogy in the regulatory elements of this pathway is usual among the Alphaproteobacteria. Although the signalling flow is usually straightforward to assess due to the configuration of the regulatory cascade (e.g. convergence from various PhyR and NepR paralogs to one EcfG σ factor (Bastiat 2010) or divergence from one PhyR-NepR stream to a number of EcfG representatives that act in series or in parallel (Lourenço *et al*., 2011; Francez-Charlot *et al*., 2016; Gottschlich *et al*., 2019), establishing functional differences between *a priori* redundant regulators may be challenging. In the case of TFA, the interplay between EcfG1 and EcfG2 in the activation of the GSR regulon had already been addressed (de Dios *et al*., 2020), depicting a model in which EcfG2 would be the master activator and EcfG1 would play an accessory role upon activation of the response, most likely as an amplifier of part of the regulon. In this work, we elucidate the signalling flow from the PhyR regulators to the EcfG σ factors via the NepR anti-σ factors based on multiple genetic, phenotypical and biochemical evidences, highlighting the specific functional differences between paralogous elements.

*In vitro* experiments addressing the interaction between the NepR1 and NepR2 anti-σ factors and the EcfG1 and EcfG2 σ factors clearly indicate that, although both proteins bind EcfG1 and EcfG2, NepR1 interacts more efficiently with EcfG1 and EcfG2 than NepR2. This is a remarkable difference with previously described NepR-EcfG interactions, such as those of *C. crescentus* (Lourenço *et al*., 2011) and *S. melonis* (Kaczmarczyk *et al*., 2011). In the first case, the main NepR element does not interact with the secondary EcfG paralog. In TFA, the *in vitro* transcription assays and interaction quantifications show that NepR1 efficiently binds both EcfG σ factors, ruling out that possibility. In the case of *S. melonis* a secondary NepR protein (also termed NepR2) is unable to be co-expressed with EcfG1 which has suggested an inefficient interaction between them (Gottschlich *et al*., 2019). In TFA, IVT assays show that NepR2, in amounts sufficiently high (10:1 molecular excess with respect to either σ factor), is able to inhibit around 75% of the transcription driven by either EcfG1 or EcfG2. This hints that the interactions between NepR2 and both EcfG1 and EcfG2 in TFA would occur mainly upon GSR activation, when the *nepR2* transcription has already been induced and the respective protein product is present in sufficiently high cellular concentrations. On the other hand, before GSR activation, inhibition by NepR2 would be less prominent due to its negligible amounts, yet essential, compared to the inhibition exerted by NepR1. The *in vitro* differences between NepR1 and NepR2 are coherent with the *in vivo* expression measurements obtained with the *nepR2::lacZ* reporter in the *nepR* mutant backgrounds, assigning to NepR1 the role of controlling the initial activation of the GSR upon stress exposure. Later on, NepR2 would act once the response is active by modulating its final intensity. This role as feedback modulator has been discussed for other additional NepR orthologs (Francez-Charlot *et al*., 2016; Gottschlich *et al*., 2019) whose mechanistic insights will be further discussed below.

When various NepR paralogs are present, they may exhibit functional differences, such as those NepR pairs characterised in *M. extorquens* and *S. melonis*. In these species, the main NepR element binds either the main EcfG σ factor or PhyR, depending on the phosphorylation state of the response regulator. Oppositely, the secondary NepR paralog (MexAM1_META2p0735 and NepR2, respectively) interacts with PhyR, but it seems unable to form a stable complex with any of the EcfG paralogs encoded in these species (Francez-Charlot *et al*., 2016; Gottschlich *et al*., 2019). This regulatory interplay supports a model in which, once the activation of the GSR is triggered by the PhyR-dependent sequestration of the main NepR, the production of a paralogous NepR would titrate PhyR in a negative feedback loop so that a proportion of the primary NepR is available to inhibit the σ factor activity. The balance in the amounts of NepR bound either to EcfG or to PhyR would determine the levels of GSR activity. Furthermore, this has been proposed as a mechanism to rapidly switch off the response when the stress disappears (Gottschlich *et al*., 2019). The IVT results obtained with the TFA regulators, together with the protein amounts of the two NepR anti-σ factors before and after triggering the response, provide direct evidence to support this indirect negative feedback regulation. Also, NepR1 binds EcfG1 and EcfG2 more efficiently than NepR2, whereas the latter is able to interact with PhyR1 and PhyR2 (in their phosphorylated state) more efficiently than NepR1. Hence, the GSR would be modulated by a two-level negative feedback loop in TFA, with NepR2 playing a dual role: i) directly inhibiting the EcfG1 and EcfG2 activity (mainly under GSR-inducing conditions and to a lesser extent in the absence of stress), and ii) indirectly inhibiting the GSR activity by titrating the active PhyR proteins (and thus releasing NepR1 to inhibit EcfG1 and EcfG2) to prevent the overactivation of the system.

Biochemical studies on the NepR-PhyR interaction (Luebke *et al*., 2018) revealed that its specificity is determined by the NepR intrinsically disordered N-terminal region, termed FR1, particularly in the residues adjacent to the helix α1. This region also participates in the PhyR activation by enhancing its phosphorylation (Kaczmarczyk *et al*., 2014; Herrou *et al*., 2015; Luebke *et al*., 2018). Also, the FR1 fragment shows a strong divergence even comparing NepR paralogs encoded within the same strain, such as the TFA NepR1-NepR2 pair and the *S. melonis* NepR-NepR2 pair (Sup. Fig. S7). In agreement with Luebke *et al*. (2018), this region, especially in the fragment right next to the α1 helix, was the most divergent between main and additional NepR paralogs, which may suggest different specificities for the respective EcfG and PhyR proteins. These observations might explain the distinct interplay between NepR1 and NepR2 and the rest of regulators in this pathway, hinting at a modulatory role of NepR2 beyond the usual σ-anti-σ titration.

Ascending further upstream in the GSR cascade, we tackled the characterisation of the two PhyR proteins encoded in TFA. In other Alphaproteobacteria with two PhyR paralogs (e.g. RsiB1 and RsiB2 from *S. meliloti* (Bastiat *et al*., 2010)), both elements appear to exert a similar control on the GSR, since their mutation led to similar phenotypes. However, in TFA both PhyR1 and PhyR2 seem to play different functional roles as judged by the stress resistance assays testing the single and double Δ*phyR* mutants. These experiments indicate a specificity in the signalling depending on the stress that triggers the response. Nevertheless, given the nature of these regulators and their role in the signalling, it seems clear that they do not participate in the specific sensing themselves. Instead, there would be other elements above the PhyR level, such as the four putative HRXXN histidine kinases predicted in the TFA genome (SGRAN_1165, SGRAN_1773, SGRAN_2544 and SGRAN_3485) or any other phosphor-transfer element yet unknown, the ones differentiating among signals and/or transducing them selectively to either PhyR1, PhyR2 or both of them. The role of each PhyR regulator in this pathway was addressed measuring the activity of the *nepR2::lacZ* reporter under carbon starvation, a condition that seems to trigger the signalling through both PhyR elements (Fig. 4), although to different extends. The differences in GSR activation *in vivo* and the ability of each PhyR protein to stimulate transcription *in vitro* indicate that PhyR1 is able to produce a stronger activation of the GSR compared to PhyR2. This would imply that PhyR1 and PhyR2 have different binding affinities for NepR1 and NepR2, eventually affecting the proportion of active EcfG σ factors, and thus, the intensity of the response. Taken together, the stress specificity showed by PhyR1 and PhyR2 and their different abilities to bind NepR1 and NepR2 would suggest a mechanism to modulate the intensity of the GSR output accordingly to the stress that triggered it.

Taking together all the results obtained throughout this work, a step-wise GSR regulatory model proposed for TFA would be as depicted in Fig. 6. When some kind of stress appears either in the environment or in the cytoplasm, it would be sensed by one or more of the predicted HRXXN histidine kinases, causing an autophosphorylation in their conserved His residue. The signal would be transduced in a specific manner, either directly or indirectly, to PhyR1 and/or PhyR2, which would receive the phosphoryl in an Asp residue. The phosphorylation would trigger a conformational change to expose their σ-like domains. This would lead to the sequestration of NepR1 in a different proportion, depending on whether the signalling occurred through PhyR1 and/or PhyR2. NepR1 titration would release EcfG2 and the basal amount of EcfG1 from inhibition, thus activating the GSR regulon. As part of that regulon, the expression of the *nepR2ecfG1* operon would be induced, increasing EcfG1 and, more importantly, NepR2 levels. In a negative feedback loop, NepR2 would inhibit EcfG1 and EcfG2 in a direct manner by protein-protein interaction. Also, NepR2 would bind PhyR1 and/or PhyR2 with higher affinity than NepR1, titrating them away from the latter. After its release, NepR1 would be again available for directly inhibiting EcfG1 and EcfG2 together with the remaining NepR2. The effect of NepR2 at the EcfG and PhyR levels, together with its high accumulation, would ensure autoregulated levels of GSR by a negative feedback loop to prevent overactivation or to quickly switch GSR off.

**Figure 6.**
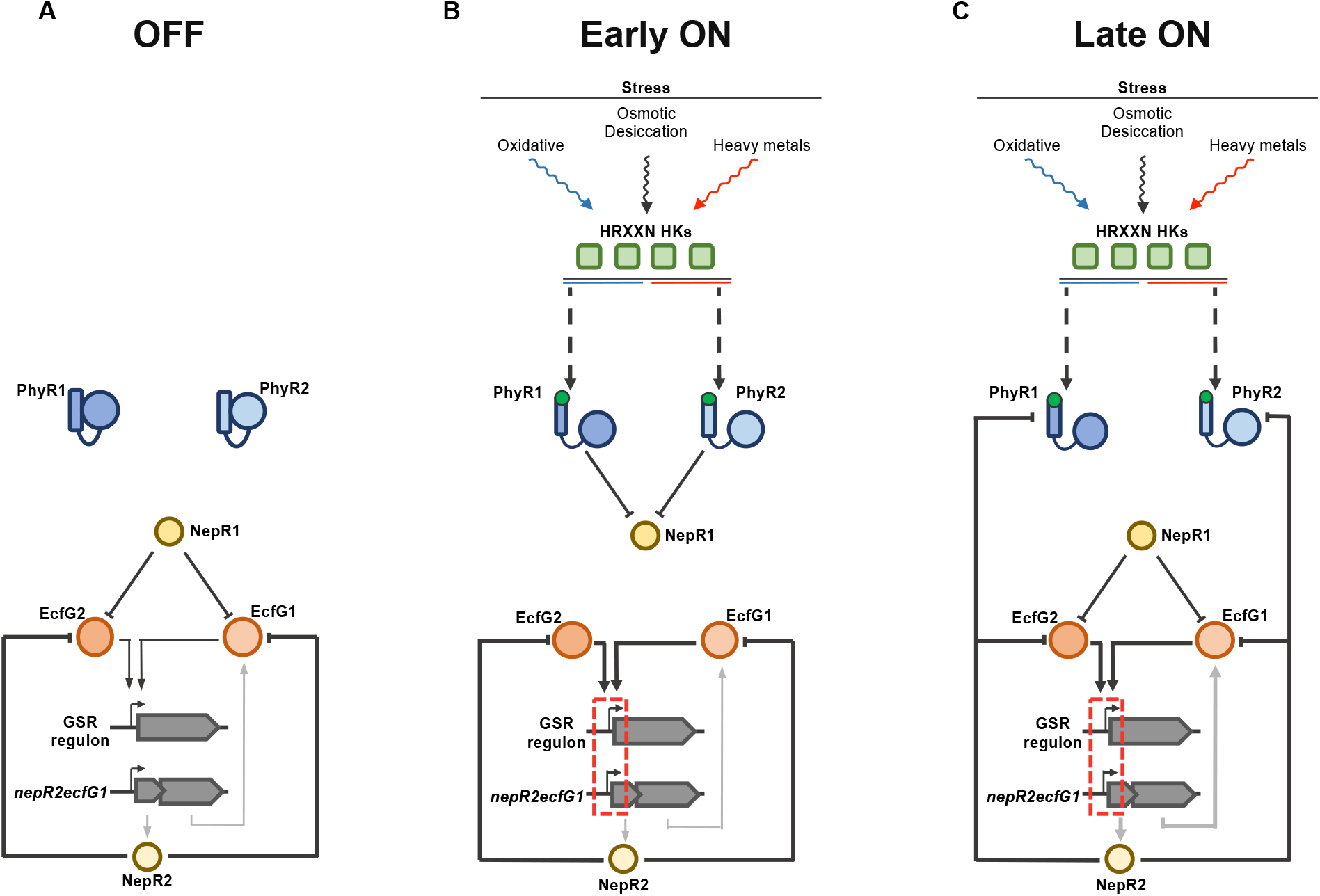
Step-wise representation of the regulatory model for the GSR signalling pathway in *S. granuli TFA*, indicating the interplay among the regulators and/or their state in the absence of stress (A), at the onset of the stress signalling (B) and once the GSR is fully active (C). Green squares represent the four histidine kinases annotated in the TFA genome, PhyR regulators are represented in dark (PhyR1) and light blue (PhyR2, NepR anti-σ factors are represented in dark (NepR1) and light yellow (NepR2), EcfG σ factors are represented in light (EcfG1) and dark orange (EcfG2), genes are represented in grey. Wavy arrows indicate stress sensing (black lines indicate signalling through PhyR1 and PhyR2, blue lines indicate signalling through PhyR1, red lines indicate signalling through PhyR2); dashed arrows indicate phosphosrylation of the PhyR regulators (either direct or through intermediate elements); a green circle represents the phosphorylation of PhyR1 and PhyR2; black arrows indicate a regulatory relationship by direct interaction (triangular arrowheads indicate a positive effect, flat arrowheads indicate a negative effect); grey arrows represent transcription and translation.

## MATERIALS AND METHODS

### Media and growth conditions

*Escherichia coli* and *Sphingopyxis granuli* strains were routinely grown in LB rich medium (Sambrook *et al*., 1989) at 37 ºC or MML mineral medium (Andujar *et al*., 2000) at 30 ºC, respectively. When indicated, *S. granuli* strains were grown in minimal medium (Dorn *et al*., 1974) supplemented with β-hydroxybutyrate (BHB) as a carbon source in concentrations 8 or 40 mM, depending on the experimental conditions. When appropriate, solid and liquid media were supplemented with kanamycin (25 mg/l for *E. coli*, 20 mg/l for *S. granuli*), ampicillin (100 mg/l for *E. coli*, 5 mg/l for *S. granuli*), streptomycin (50 mg/l for routine selection, 200 mg/l for selection of co-integrates of pMPO1412-derivative plasmids) or X-gal (25 mg/l).

### Plasmids, strains and oligonucleotides

Bacterial strains, plasmids and oligonucleotides used in this work are indicated in Sup. Table S1.

For the generation of mutant strains with scar-less chromosomal modifications (deletions/insertions), the SceI double-strand break mediated double recombination procedure was followed as previously described (González-Flores *et al*., 2019; de Dios *et al*., 2020). Briefly, a pMPO1412-derivative plasmid containing upstream and downstream 1 kb flanking regions of the fragment to be deleted or the position where the insertion will be placed was transformed in the respective *S. granuli* parental strain and its recombination into the chromosome was selected in the presence of kanamycin. Double-check of recombinant candidates or co-integrates was performed by growing them in the presence of streptomycin (200 mg/l). Subsequently, plasmid pSWI (Martinez-Garcia & de Lorenzo, 2011), bearing the SceI open reading frame, was transformed into the co-integrate strain to force a second recombination event. Candidates bearing the desired modifications were checked by PCR. For the construction of strains with multiple modifications, this procedure was performed serially with the different pMPO1412-derivative plasmids. Deletion mutants constructed with this strategy were MPO865 (with pMPO1416), MPO866 (with pMPO1414), MPO867 (with pMPO1415), MPO868 (with pMPO1414 and pMPO1415), MPO889 (with pMPO1416 using MPO860 as parental strain), MPO898 (with pMPO1428) and MPO899 (with pMPO1428 using MPO865 as parental strain). Strains bearing 3xFLAG-tagged genes constructed following this protocol were MPO906 (with pMPO1453), MPO907 (with pMPO1454), MPO908 (with pMPO1457) and MPO909 (with pMPO1458).

Strains bearing the *nepR2::lacZ* reporter inserted in the chromosome (MPO871, MPO872, MPO873, MPO874, MPO890, MPO900 and MPO902) were constructed by transforming the respective parental strain with plasmid pMPO1408 and selecting its chromosomal integration by a single recombination event.

For construction of pMPO1412 derivatives, the respective upstream and downstream flanking regions were amplified using *S. granuli* TFA genomic DNA as template and were subsequently assembled together by overlapping PCR. Oligonucleotide pairs used in each case were phyR1 del1 SacI-phyR1 del2 and phyR1 del3-phyR1 del4 BamHI for pMPO1414; phyR2 del1 BamHI-phyR2 del2 and phyR2 del3-phyR2 del4 EcoRI for pMPO1415; nepR1 del1 SacI-nepR1 del2 and nepR1 del3-nepR1 del4 BamHI for pMPO1416; ecfG1 FLAG-1 BamHI-ecfG1 FLAG-2 and ecfG1 FLAG-3-ecfG1 FLAG-4 SacI for pMPO1453; ecfG2 FLAG-1 BamHI-ecfG2 FLAG-2 and ecfG2 FLAG-3-ecfG2 FLAG-4 EcoRI for pMPO1454; nepR1-FLAG-1-nepR1-FLAG-2 and nepR1-FLAG-3-nepR1-FLAG-4 for pMPO1457; nepR2-FLAG-1-nepR2-FLAG-2 and nepR2-FLAG-3-nepR2-FLAG-4 for pMPO1458. For construction of pMPO1414, pMPO1415, pMPO1416, pMPO1453 and pMPO1454, the assembled PCR fragments were digested with the appropriate restriction enzymes (included in the name of the respective oligonucleotides) and ligated into pMPO1412 digested with the same enzymes. In the case of pMPO1457, pMPO1458, pMPO1459 and pMPO1460, the assembled PCR fragments were directly cloned into pMPO1412 cut with SmaI.

The pMPO1412 derivative pMPO1428, used for the deletion of the *nepR2ecfG1* operon, was constructed based on the previously constructed pMPO1407 and pMPO1413 (de Dios *et al*., 2020). pMPO1407 was digested with XhoI, blunted with Klenow and subsequently cut with Acc65I. The resulting 1 kb fragment was ligated into pMPO1413 digested with StuI and Acc65I.

pTXB1 and pTYB21 derivatives for protein overproduction were constructed based on the guidelines provided with the IMPACT kit (New England Biolabs). The coding sequences of *nepR1, nepR2, phyR1* and *phyR2* were amplified by PCR from *S. granuli* TFA genomic DNA using oligonucleotide pairs ORF-nepR1 fw-ORF-nepR1 rv BamHI, ORF-nepR2 fw-ORF-nepR2 rv BamHI, ORF-phyR1 fw NdeI-ORF-phyR1 rv and ORF-phyR2 fw NdeI-ORF-phyR2 rv, respectively. *nepR1* and *nepR2* fragments were digested with BamHI and ligated into pTYB21 cut with SapI, blunted with Klenow and digested with BamHI, resulting in plasmids pMPO1434 and pMPO1435, respectively. *phyR1* and *phyR2* fragments were digested with NdeI and ligated into pTXB1 cut with SapI, blunted with Klenow and digested with NdeI, resulting in plasmids pMPO1436 and pMPO1437, respectively.

### Stress phenotypic assays

Stress resistance assays were performed as in de Dios *et al*. (2020). Briefly, to test the resistance to osmotic stress and copper, 10 μl spots of serial dilutions of late-exponential phase cultures were placed on solid MML rich medium plates supplemented with NaCl 0.6 M or CuSO_4_ 3.5 mM and incubated for 5 days at 30 ºC. For desiccation assays, 5 μl spots of serial dilutions of late-exponential phase cultures were placed on 0.45 μm pore size filters (Sartorius Stedim Biotech GmbH) and they were left to air-dry in a laminar flow cabin for 5 h (5 min in the control assay). Then, filters were placed on MML rich medium plates supplemented with bromophenol blue 0.002% and incubated for 5 days at 30 ºC. In the case of recovery from oxidative shock, late-exponential phase cultures were diluted to an OD_600_ of 0.1 in MML medium. When an OD_600_ 0.5 was reached, H_2_O_2_ was added to the medium in a final concentration of 10 mM. Recovery from the treatment is represented by a percentage of the OD_600_ reached by treated cultures after 5 h of growth compared to non-treated cultures. At least three independent replicates of each experiment were performed, and most representative examples are shown.

### GSR activation assays and expression measurements

Saturated preinocula were diluted to an OD_600_ of 0.05 in minimal medium supplemented with β-hydroxybutyrate 40 mM and incubated at 30 ºC in an orbital shaker for 16 h. Then, 20 ml of minimal medium with β-hydroxybutyrate 8 mM were inoculated at OD_600_ 0.1. β-galactosidase activity (Miller, 1972) from the *nepR2::lacZ* reporter was measured after 10 h and 58 h of growth, representing exponential and stationary phase, respectively (Sup. Fig. 3). Averages of three independent replicates are represented.

### Protein overexpression and purification

*S. granuli* TFA core RNA polymerase, EcfG1 and EcfG2 were purified as previously published in de Dios *et al*. (2020).

NepR1, NepR2, PhyR1 and PhyR2 proteins were overexpressed and purified using the IMPACT kit (New England Biolabs) following the manufacturer’s instructions and equal procedures for the four of them. Briefly, pMPO1434, pMPO1435, pMPO1436 and pMPO1437 (for overexpression of *nepR1, nepR2, phyR1* and *phyR2*, respectively) were transformed into *E. coli* ER2566 host strain. Saturated pre-inocula of each plasmid-bearing strain were diluted to an OD_600_ of 0.1 in different total volumes of LB medium, depending on the gene to be overexpressed (2 l for *nepR1*, 1 l for *nepR2*, 4 l for *phyR1* and 1 l for *phyR2*), and incubated at 37 ºC in an orbital shaker until reaching OD_600_ 0.7. Then, cultures were chilled on ice and subsequently induced with IPTG 0.5 mM and incubated overnight in a shaker at 16 ºC. After harvesting the cultures and assessing the induction by SDS-PAGE, cell pellets were resuspended in binding buffer (Tris-HCl 20 mM pH 8, NaCl 0.5 M), lysed by sonication and clarified by centrifugation. Once the chitin resin was packed in a purification column and washed with binding buffer, the respective clarified lysates were loaded on the column and left to flow through the resin by gravity at a low flow rate. Afterwards, the column was flushed with 100 ml of binding buffer prior to the induction of the on-column protein cleavage. To release the target protein, the resin was incubated with TEDG buffer (Tris-HCl 50 mM pH 8, glycerol 10%, Triton X-100 0.01%, EDTA 0.1 mM, NaCl 50 mM) supplemented with DTT 50 mM at 18 ºC for 40 h approximately. The eluate content in the target protein was assessed by SDS-PAGE. Then, DTT concentration in the buffer was reduced by dialysis against TEDG buffer with DTT 0.1 mM at 4 ºC overnight using a 3 KDa pore size dialysis cassette (ThermoFisher Scientific). Finally, purity and concentration of the protein mixtures were evaluated by densitometry comparing to different dilutions of BSA using a Typhoon scanner and the ImageLab software. For long-term storage, protein mixtures were aliquoted and frozen at -80 ºC.

#### *In vitro* transcription

Multi-round *in vitro* transcription (IVT) reactions were performed as in Porrua *et al*. (2009) with modifications from de Dios *et al*. (2020). Briefly, reactions were run in a final volume of 22.5 μl in IVT buffer (Tris-HCl 10 mM pH 8, NaCl 50 mM, MgCl_2_ 5 mM, KCl 100 mM, BSA 0.2 mg/ml, DTT 2 μM) at 30 ºC. A mixture containing the appropriate combination of the different GSR regulators, either supplemented or not with acetyl phosphate 15 mM depending on the experiment, was preincubated at 30 ºC for 5 min. In this mixture, to ensure that any transcriptional activation would be due to disruption of the EcfG-NepR interaction by PhyR, the right amount of each PhyR protein (with or without acetyl phosphate) was added first in a tube chilled on ice followed by a volume bearing the appropriate EcfG-NepR pair also pre-incubated on ice. 0.2 μM of the respective EcfG σ factor was set as reference to stablish molecular proportions with the rest of the regulators present in the reaction. After that, the core RNA polymerase mix was added to the reaction and it was incubated for 5 min. Subsequently, 0.5 μg of plasmid pMPO1440 were added as circular template. 10 min later, a mix of ATP, GTP, CTP (final concentration of 0.4 mM), UTP (0.07 mM) and [α-32P]-UTP (0.33 mM, Perkin Elmer) was added to start the reaction. After 10 min, reaction re-initiation was prevented by adding heparin to a final concentration of 0.1 mg/ml, and 10 min later reactions were arrested by adding 5 μl of stop/loading buffer (0.5 % formamide, 20 mM EDTA, 0.05% bromophenol blue, 0.05% xylene cyanol). Samples were boiled for 3 min and run in a 4% polyacrylamide-urea denaturing gel in TBE buffer at room temperature. Gels were dried and exposed in a phosphoscreen and results were visualised in an Amersham Typhoon scanner and analysed using the ImageQuant software (both provided by GE Healthcare Bio-Sciences AB). Quantifications refer to the median intensity of each band normalised against the levels of transcription obtained by each EcfG protein alone, in the absence of PhyR and NepR. The figure shows a representative assay of this experiment and quantifications are the average of three independent replicates.

### Protein immunodetection (Western blot)

Samples were obtained from cultures in exponential and stationary phase as explained above for gene expression assays. For each sample,1 OD_600_ unit was harvested by centrifugation and the cell pellet was resuspended un 25 μl bidistilled water. Whole-cell protein content was measured using the RC DC Protein Assay kit (Bio-Rad) and the remaining sample was mixed with loading buffer 2X, boiled for 5 min and centrifuged. The equivalent volume to 10 μg of protein was run in a Stain-Free FastCast 12.5% polyacrylamide gel (Bio-Rad) and transferred to a nitrocellulose membrane using the Trans-Blot Turbo semi-dry system (Bio-Rad) following the manufacturer’s instructions. The membrane was washed with TTBS buffer and blocked with 5% skimmed milk powder in TTBS buffer (blocking solution). Subsequently, the membrane was incubated overnight with a 1:2000 dilution of mouse monoclonal anti-FLAG antibody (Sigma-Aldrich) in blocking solution at 4 ºC with mild shaking. Then, the membrane was washed with TTBS, incubated for 2 h with a 1:10000 dilution of anti-mouse secondary antibody (Sigma-Aldrich) in blocking solution at room temperature with mild shaking and washed again with TTBS. Finally, the membrane was developed with the Immun-Star AP Chemiluminescence kit (Bio-Rad) and the signal was detected with a ChemiDoc image system (Bio-Rad) and analysed with the ImageLab software (Bio-Rad). Representative experiments from three independent replicates are shown. Quantifications refer to the average fold-change in stationary phase compared to exponential phase of three independent replicates.

### Surface plasmon resonance

EcfG1 and EcfG2 interaction kinetics with respect to immobilised NepR1 or NepR2 were measured using a BIAcore X100 device (GE Healthcare Life Sciences). Assays were performed at 30 ºC in TEDG buffer. NepR1 (12.1 RU) or NepR2 (36.4 RU) were immobilised in on the surface of a CM5 chip using 10 mM acetate buffer pH 4.0 or 5 mM malate buffer pH 5.5, respectively, at 30 ºC with a contact time of 300 s, following the manufacturer’s instructions. Serial 2-fold dilutions of EcfG1 and EcfG2 in TEDG buffer were injected in the system at a flow rate of 20 μl/min in concentrations ranging from 60 nM to 0.469 nM. Analyte contact time was enough to reach interaction equilibrium and dissociation time was 300 s. After each interaction cycle, the chip was regenerated by injection of 10 mM glycine-HCl buffer pH 2.0. Data were fitted to a 1:1 interaction model using the evaluation software provided by the manufacturer (GE Healthcare Life Sciences). Reliability of the results was assessed according to U-value < 15 and χ^2^ < 5%R_max_. Interaction affinity was defined by the dissociation constant (K_D_) obtained for each NepR-EcfG pair. At least three independent replicates were assayed for each pair.

## Acknowledgements

We wish to thank Guadalupe Martín and Nuria Pérez for technical help and all members of the laboratory for their insights and helpful suggestions. This work was supported by grant BIO2014–57545-R, co-funded by the Spanish Ministry for Education and Science and the European Regional Development Fund, and by the FPU fellowship (Ref. FPU15/04789, Ministerio de Educación y Ciencia, Spain), awarded to R.D.

## Author contributions

R.D. performed the experiments, R.D., F.R.-R. and E.S. designed the experimental strategy and analysed the results. F.R.-R. and E.S. supervised the work. R.D. and F.R.-R. wrote the manuscript considering the revisions of all the authors.

